# Femtosecond laser preparation of resin embedded samples for correlative microscopy workflows in life sciences

**DOI:** 10.1101/2023.01.10.523473

**Authors:** C Bosch, J Lindenau, A Pacureanu, CJ Peddie, M Majkut, AC Douglas, R Carzaniga, A Rack, L Collinson, AT Schaefer, H Stegmann

**Author notes:** Authors contributed equally (C Bosch, J Lindenau).

## Abstract

Correlative multimodal imaging is a useful approach to investigate complex structural relations in life sciences across multiple scales. For these experiments, sample preparation workflows that are compatible with multiple imaging techniques must be established. In one such implementation, a fluorescently-labelled region of interest in a biological soft tissue sample can be imaged with light microscopy before staining the specimen with heavy metals, enabling follow-up higher resolution structural imaging at the targeted location, bringing context where it is required. Alternatively, or in addition to fluorescence imaging, other microscopy methods such as synchrotron X-ray computed tomography with propagation-based phase contrast (SXRT) or serial blockface scanning electron microscopy (SBF-SEM) might also be applied. When combining imaging techniques across scales, it is common that a volumetric region of interest (ROI) needs to be carved from the total sample volume before high resolution imaging with a subsequent technique can be performed. In these situations, the overall success of the correlative workflow depends on the precise targeting of the ROI and the trimming of the sample down to a suitable dimension and geometry for downstream imaging.

Here we showcase the utility of a novel femtosecond laser device to prepare microscopic samples (1) of an optimised geometry for synchrotron X-ray microscopy as well as (2) for subsequent volume electron microscopy applications, embedded in a wider correlative multimodal imaging workflow (**Fig. 1**).

## INTRODUCTION AND RESULTS

### Femtosecond laser milling of resin-embedded specimens

A femtosecond laser (fs laser) is an ultra-short pulse laser with pulse lengths typically ranging between a few ten and a few hundred femtoseconds. A fs laser allows rapid material removal to access targeted regions within a larger sample. Here, we used a fs laser with a wavelength of 515 nm, pulse length <350 fs, adjustable pulse repetition rate of 1 kHz to 1 MHz and focus spot diameter of <15 µm. The laser was integrated in a FIB-SEM system (Zeiss Crossbeam 350) allowing for a rapid sample and laser target coordinate transfer between the laser processing chamber and the main chamber of the instrument (**Fig. 2**). The fs laser was positioned in a separate chamber attached directly to the main Crossbeam chamber via an airlock. This design allows the ablation of large sample volumes without contaminating detectors or other sensitive components of the main chamber. Since fs lasers provide athermal material ablation ^1^, the laser affected zone (LAZ) is minimised, extending less than 0.5 µm into a laser cut cross-section. The narrow LAZ enables the targeting of milling boundaries with an accuracy in the range of µm. The geometry of the carved specimen can be arbitrarily defined by the operator, including curved surfaces, making it trivial to generate samples in the shape of cylindrical pillars, a feature also at reach when milling the samples using lathe systems ^2^ but not when using microtome-based approaches. Fs lasers enable high ablation rates of up to several 10^6^ µm^3^/s, depending on sample material and the laser settings used, and are compatible with heavy metal-stained, resin-embedded biological soft tissues ^3,4^.

**Figure 1.**
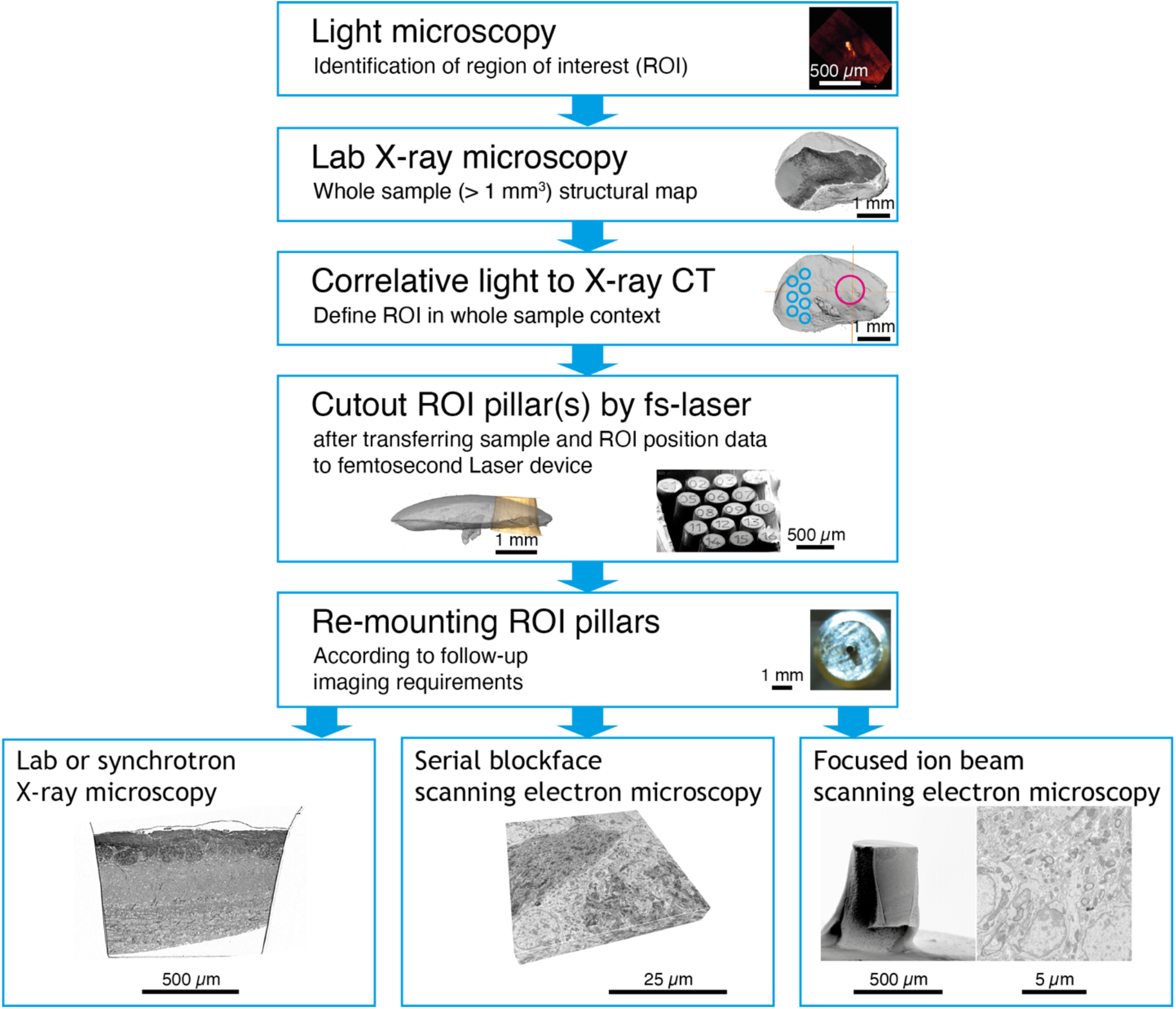
Correlative microscopy workflows incorporating samples processed using a femtosecond laser.

**Figure 2.**
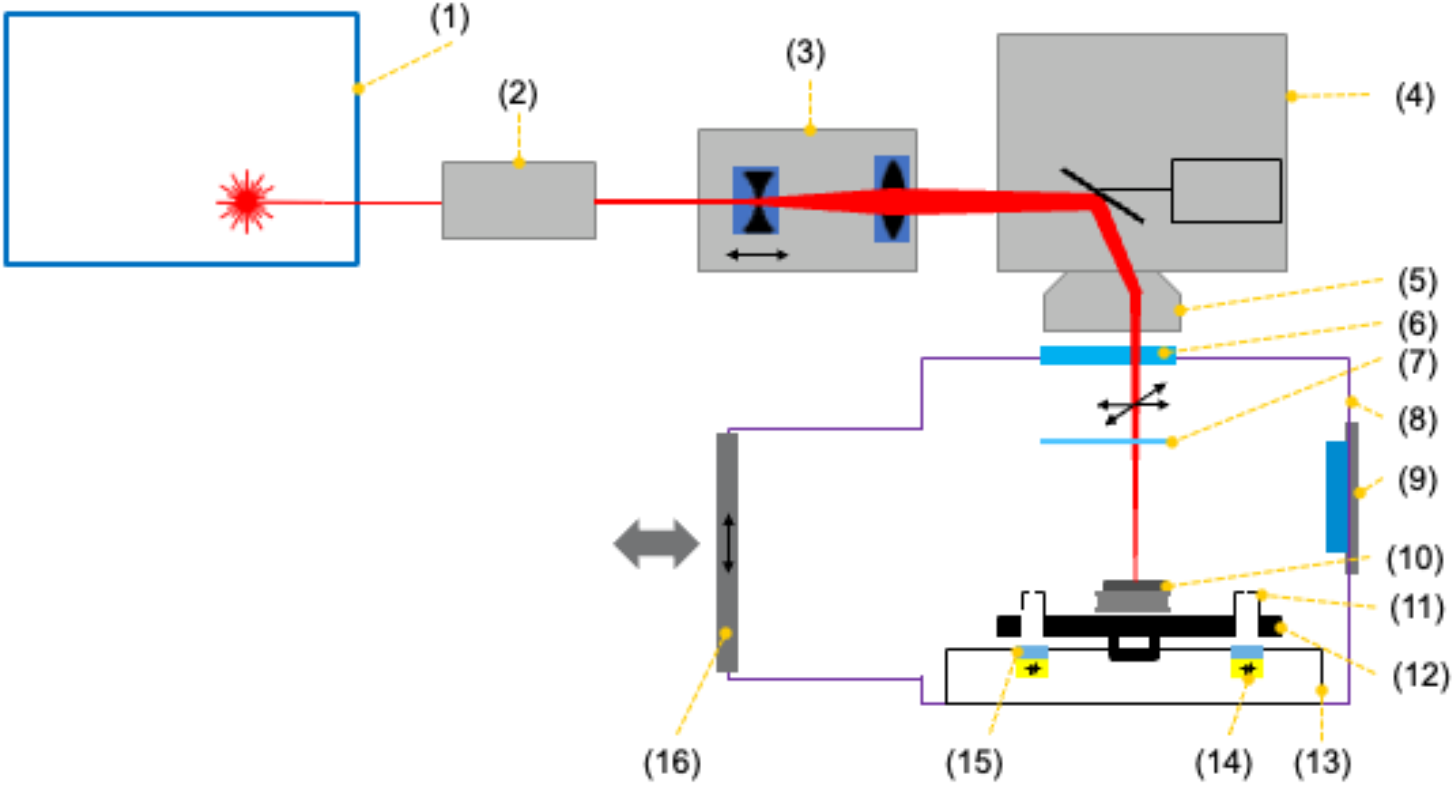
Scheme of a femtosecond laser sample milling device. (1) femtosecond (fs) laser source; (2) beam expander; (3) defocus unit (Z-axis scanner); (4) laser scanning unit; (5) telecentric f-theta objective; (6) laser entry window (anti-reflective coated); (7) protective window (no anti-reflective coating, replaceable); (8) laser preparation chamber; (9) observation window (covered by movable slider); (10) sample mounted on stub; (11) registration apertures; (12) laser specimen holder; (13) specimen holder mount; (14) diode-based registration unit; (15) attenuation filter disk; (16) airlock to main crossbeam chamber.

Altogether, these characteristics make the fs laser a suitable tool to extract targeted volumes of interest from a larger sample, the size, coordinates, and geometry of which were previously defined with compatible imaging techniques. Moreover, since the milling operation only requires access to one face of the resin block, multiple milling operations can be performed on a single specimen, enabling the extraction of tens of samples from neighbouring regions in the same specimen.

This versatility places fs lasers in a potentially highly useful niche, enabling the generation of multiple independent specimens containing targeted volumes of interest from sub-mm^3^ to multiple mm^3^ in size, and with optimised geometry for follow-up imaging techniques. In these specimens, the histological detail can be efficiently retrieved at multiple scales in 3D using several distinct imaging technologies – from synchrotron X-ray computed tomography with propagation-based phase contrast ^5^ or nano-holographic tomography ^6^, to serial blockface SEM ^7-9^, and ultimately FIB-SEM ^10,11^.

### Sample trimming in biomedical sample preparation protocols

The use of fs lasers for targeted sample preparation in materials science is a well-established method and has been applied in a variety of different studies ^12,13^. Utilisation of the fs laser for sample processing in life sciences provides a tool for targeted trimming of soft biological tissue that can complement other available mechanical tools in workflows that are highly dependent on precise specimen trimming, such as ultramicrotomy ^14^ or lathe implementations ^2^. To interrogate the ultrastructure at nanometre scale, tissue specimens of dimensions reaching several cubic millimetres need to be contrasted with heavy metals and embedded in resin ^15-22^. Studying rare, localised structures with sub-µm detail requires a highly accurate preparation of those volumes of interest to be imaged and analysed ^20,23-25^. Both preparation and imaging techniques will often impose sample size restrictions. Altogether, experimental targeting and sample size restrictions will define how the stained specimen will have to be trimmed, to ultimately match the downstream imaging modality. Finally, extracting multiple targeted samples from a single stained specimen is very challenging with conventional techniques, and therefore at this point it is more common to aim to prepare only one final sample per resin-embedded specimen. This, however, not only limits the useability of potentially highly precious specimens, but also prevents the combined analysis of spatially distant ultrastructural features.

When looking at X-ray microscopy (XRM) of resin-embedded biological samples, additional challenges and limitations apply for sample preparation. In this case, for optimal results, samples are preferred to be cylindrical in geometry to reduce missing edge artefacts ^26^, with specific imaging techniques and specimens imposing upper bounds on the optimal diameter range ^5^. Femtosecond laser trimming is a technique uniquely suited to prepare multiple cylindrical samples of specific diameters (200-1000 µm), all originating from a single specimen.

### Femtosecond laser for XRM and EM sample preparation

In order to assess the feasibility of a fs laser for sample preparation of biological tissue we investigated two use cases, preparation for (1) targeted X-ray computed tomography and (2) multiple samples for serial blockface or focussed ion beam volume EM acquisition.

We firstly assessed the capability of fs lasers to ablate resin (specifically epoxy resin EMbed 812 as typically used for embedding samples for electron microscopy, in particular for volume EM methods where physical sectioning is involved). Laser power, pulse frequency, scan speed and scan strategy were varied to identify settings that provided cleanly cut pillars with minimum surface roughness. With the optimum laser settings found (see methods), arrays of pillars of 450 to 830 µm height and a pillar diameter down to 250 µm (measured at the tip) were obtained.

In order to define a specific target region we employed *ex vivo* two-photon imaging in a mouse line where one specific biological structure, an olfactory bulb glomerulus ^5,27^ was genetically labelled (**Fig. 3a**). In this case, 2-photon imaging was performed after dissection of the brain, preparation of a 600 µm horizontal brain slice and GA/PFA fixation (see methods). Subsequently and in preparation for high-resolution X-ray tomography and EM, we stained the tissue with heavy metals (see methods and ^28^). Using lab X-ray tomography as described before ^5^ allowed us to identify the very region previously imaged using 2-photon microscopy (**Fig. 3b,c**). We then transferred the fixed, stained and embedded brain slice into an SEM with a coupled fs laser (**Fig. 2**) (combined FIB-SEM system, see methods, but the volume EM capabilities of this system were not used in this study). The volume containing the region of interest was delineated (magenta, **Fig. 3d**) as well as several smaller, non-targeted regions (blue). Successful targeting and separation is shown by secondary electron imaging in **Fig. 3e-f**. These cylindrical pillars could then be further imaged using multiple techniques, such as EM or synchrotron X-ray computed tomography with propagation-based phase contrast (SXRT) ^5^. Overall, milling time was 15 to 24 min to extract ∼15 pillars from a specimen. This demonstrates the possibility of using fs lasers to define features and extract ROIs in fixed, metal stained, resin-embedded biological tissue. To apply this for preparation for synchrotron X-ray tomography, we sought to obtain near-cylindrical samples with smooth surfaces as these result in the highest resolution and most-efficient data acquisition for tomography. **Fig. 3g-h** shows such an example where again a region containing a genetically labelled glomerulus is marked on a LXRT tomogram and subsequently milled on the above fs-SEM system. The resulting trimmed volume can be readily imaged at a parallel beam SXRT beamline and it contains the target region of interest, resulting in high-resolution tomograms where individual dendrites of experimentally-relevant neuronal circuits are readily visible as expected (**Fig. 3h-i**). Thus, fs lasers, targeted to specific sample regions based on XRM coordinates, can be used to reliably prepare samples optimised for X-ray tomography.

**Figure 3.**
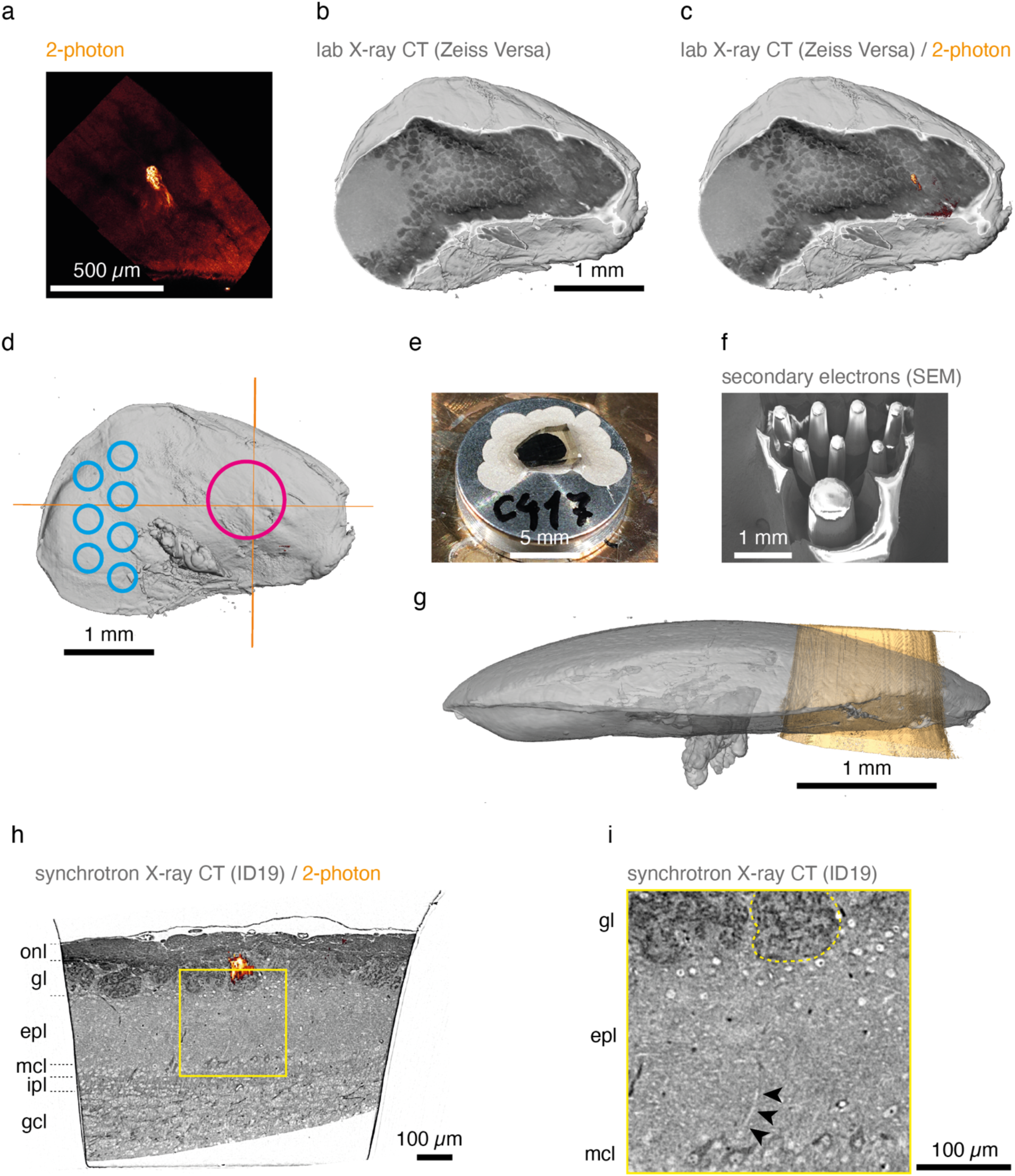
Preparation of targeted cylindrical sample for synchrotron X-ray imaging. **(a-c)** Mouse brain olfactory bulb tissue with a fluorescently labelled MOR174/9 glomerulus was imaged *ex vivo* with 2-photon microscopy, stained, embedded, and imaged with lab X-ray CT (LXRT). Both 2-photon **(a)** and LXRT **(b)** datasets were warped to the same space making it possible to identify the MOR174/9 glomerulus in the stained and embedded sample **(c). (d)** A targeted region of interest (magenta circle) was defined so it would contain the MOR174/9 glomerulus (crosshairs) and their associated projection neurons. Other non-targeted regions of interest could be also programmed in the remaining specimen (blue circles). **(e-g)** The sample was then mounted on a standard SEM pin **(e)** and the excess tissue was milled with the fs laser, leaving the carved pillars **(f)**. Finally, the targeted pillar was mounted individually and imaged with parallel-beam synchrotron X-ray microscopy with propagation-based phase contrast at the ID19 beamline of the European Synchrotron Radiation Facility **(h-i)**. The region carved corresponded to the targeted volume **(g)**.

In multiscale and multimodal imaging, high-resolution subvolumes are embedded in a lower resolution context. This is achieved by firstly performing lower resolution imaging (light microscopy or e.g. parallel beam synchrotron tomography) followed by targeted higher-resolution imaging (e.g. volume EM). Typically, a destructive sample preparation approach based on ablating or milling away the tissue is used to obtain a volume small enough for the high-resolution imaging techniques. This, however, implies that only one subvolume can be acquired at the highest resolution. For high-value specimens (e.g. those carrying lengthy functional imaging or behavioural experiments preceding anatomical investigation) it would be advantageous to acquire several neighbouring subvolumes, for example to increase the number of sample replicates. Thus, we wanted to assess whether fs laser milling could be used to create several different targeted subvolumes from one single tissue specimen. To achieve this, we employed another fixed and stained olfactory bulb slice, mechanically trimmed to a ∼(1 mm)^3^ cube (**Fig. 4a**). Using the combined fs-SEM system we marked 16 locations for individual pillars of 500 µm height and 150 µm diameter (**Fig. 4b**). To maximise the usable volume, we aimed to create pillars in close spatial proximity. Pilot experiments suggested that pillar walls could be milled as vertical as 6.0° ± 0.8° (n = 3 specimens, n’ = [13, 18, 16] pillars per specimen) (**Fig. 5a,b**). Indeed, this allowed us to reliably ablate material between marked circular regions (**Fig. 4c,d**) and extract pillars of 500 µm in height (**Fig. 5d**). This acute milling angle enabled packing pillars leaving < 200 µm of free (ablated) space between the top blockface, which makes it therefore possible to extract multiple pillars of 500 µm in height from samples with a footprint of ∼1 mm^2^ (**Fig. 5c,e-g**). Notably, not only was the entire length of the pillars freely accessible (**Fig. 4e**), but the round pillar profile was consistently smooth (**Fig. 4f**), important for many X-ray applications ^6^. After defining the target regions, milling time to cut the complete set of pillars from the specimen was ∼15 min.

**Figure 4.**
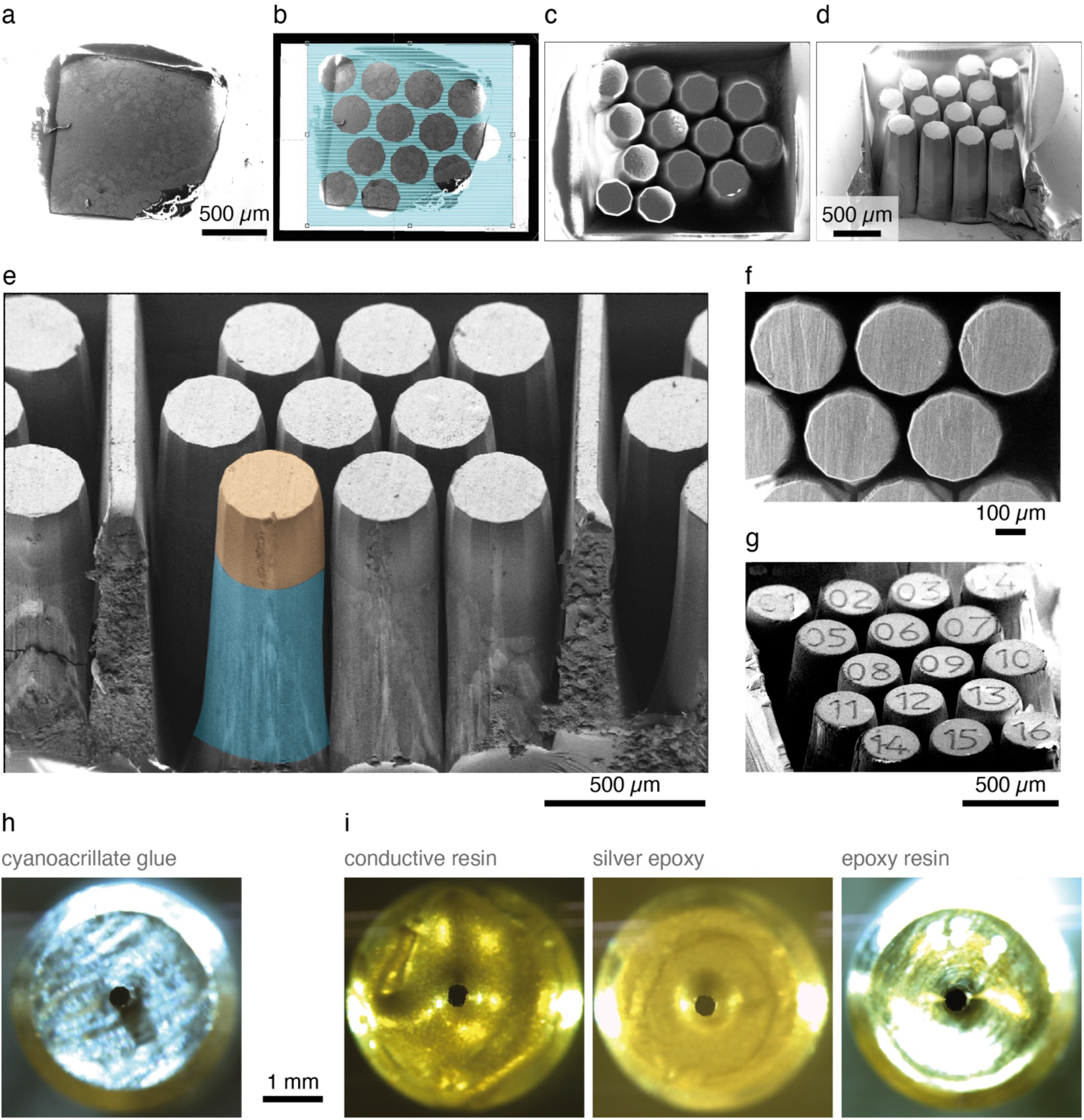
Production of pillar arrays. Multiple pillars can be extracted from a sample blockface of ∼1 mm^2^. The process involves imaging the blockface with scanning electron microscopy **(a)** and configuring the desired milling plan **(b)**. The pillars milled present sharp edges and can therefore be densely packed **(c-f)**. Finally, engraving of pillar IDs is possible following a similar procedure **(g). (h-i)** Pillars can then be separated and individually mounted **(h)** and embedded with appropriate materials to make them suitable for the intended follow-up imaging technique **(i)**.

**Figure 5.**
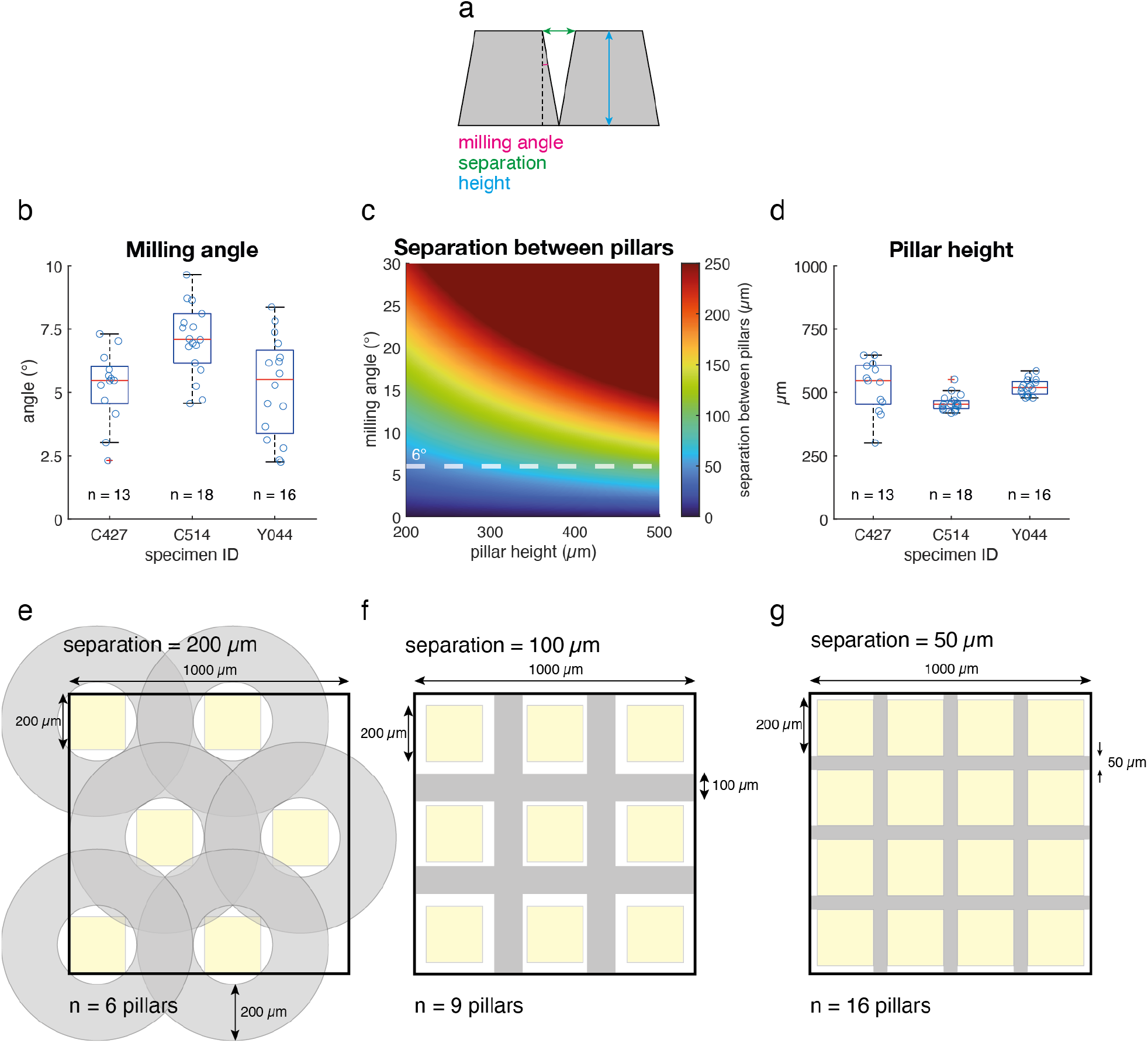
Multiple pillars can be extracted from neighbouring locations within a 1*1 mm^2^ blockface. **(a-d)** The milling angle achieved in multiple neighbouring samples enabled dense packing of pillar footprints, leaving <200 µm between the blockfaces at the sample surface, yet still consistently recovering pillars 500 µm tall. **(e-g)** Further improvements in the steepness of the milling process might allow for even tighter ROI packing density in a 1 mm^2^ blockface.

In a final step, pillars need to be separated from their common base to allow for individual re-mounting. To simplify the identification of individual pillars, we decided to “engrave” numbers to each individual pillar using the fs laser, using the same parameter settings as for the pillar milling, but directing the laser beam over the pillar top in a single pass only to create numbers with a depth of approximately 5 µm (**Fig. 4g**). This made it possible to mechanically separate all pillars simultaneously and store the pillars in individual holders for future use (**Fig. 4h-i**).

Thus, as the laser beam can be guided on any arbitrary trajectory on the sample surface, it indeed allows users to cut out any shape from a specimen and to optimise the sample geometry to the needs of the final imaging or sectioning device. It is therefore possible to prepare several samples from the initial specimen, thus increasing the experiment’s efficiency and allowing for the combined analysis of multiple spatially disperse regions in the same specimen.

## CONCLUSION

Multiscale and multimodal imaging is a very powerful approach to link the different properties and length scales typical of biological tissues. However, sample preparation presents a critical challenge. Here we introduce the fs laser as a powerful tool that complements mechanical trimming approaches. Using different brain tissue samples, we demonstrate that a fs laser allows for reliable targeting of subregions of different sizes. This is particularly pertinent when small volumes need to be extracted and separated individually for downstream imaging (e.g. for lengthy higher-resolution volume EM acquisitions) and which might require <100 µm-precise targeting. Fs lasers can be programmed to specific sample positions using coordinates from upstream volume XRM imaging (mirroring recent approaches combining XRM with automated ultramicrotome milling ^14^, albeit at significantly higher packing density (**Fig. 5**)). Importantly, sample geometry can be essentially freely chosen, making it possible to create e.g. cylindrical shapes optimised for tomography approaches.

Unlike mechanical trimming that relies on a flat blade trimming 2D faces, laser ablation operates from one single direction, ablating tissue in a line. This makes it possible to extract multiple target sample regions from one tissue specimen, including regions that are separated by only ∼100 µm (**Fig. 4**). Such a dense array of volumetric ROIs (in this case pillars) minimises any loss of material during the cutting process and enables parallel downstream processing and high resolution imaging of samples from neighbouring regions. While this packing density may not be sufficient to follow individual biological features across neighbouring pillars on their own, these can be mapped to previously obtained SXRT datasets in a correlative multimodal imaging workflow, thereby enabling inter-pillar matching of biological features at the µm scale (**Fig. 3**).

Overall, a fs laser makes it possible to reliably extract multiple samples at targeted neighbouring locations from resin-embedded soft tissue specimens and represents a very useful tool for developing robust correlative multimodal imaging workflows in life sciences.

## Acknowledgements

This research was funded in whole, or in part, by the Wellcome Trust (FC001153). For the purpose of Open Access, the author has applied a CC BY public copyright licence to any Author Accepted Manuscript version arising from this submission. This work was carried out with the support of Diamond Light Source, beamline I13-2 (proposal 20274). This work was supported by the Francis Crick Institute, which receives its core funding from Cancer Research UK (FC001153), the UK Medical Research Council (FC001153), and the Wellcome Trust (FC001153), and by a Physics of Life grant to ATS and AP (grant reference EP/W024292/1). AP has received funding from the European Research Council under the European Union’s Horizon 2020 Research and Innovation Programme (grant no. 852455). ATS was a Wellcome Trust Investigator when data was acquired and analysed (110174/Z/15/Z).

We are grateful to the biological research, electron microscopy, and scientific computing science technology platforms of the Francis Crick Institute and additionally thank Anne Bonnin, Catherine MacLachlan and members of the Sensory Circuits and Neurotechnology Lab for discussion.

## Author contributions

CB, JL, ACD, RC, ATS and HS conceptualised and CB, JL, ATS and HS administered the project. CB, AP, CJP, MM, AR and HS were involved in investigation, methodology and data curation, CB and HS in software, formal analysis and validation and CB, JL and HS in visualisation. JL, AP, ACD, RC, AR, LC, ATS and HS provided funding and resources, and AR, LC, ATS and HS supervised and guided research. CB, JL, ATS, HS wrote the original draft and all authors provided input, reviewed and edited the manuscript.

## Declaration of Interest

JL, ACD and HS are employees of Carl Zeiss Microscopy GmbH, the manufacturer of the femtosecond laser and of the Zeiss Crossbeam FIB-SEM employed and evaluated in this study. The authors have no further relevant interests to disclose.

## Materials and Methods

### Animals

All animals used were 8–24 week-old mice of C57/Bl background and mixed gender. For Figure 3 we used a male transgenic mouse with a genetically labelled glomerulus MOR174/9-eGFP as in ^27^. All animal protocols were approved by the ethics committee of the board of the Francis Crick Institute and the United Kingdom Home Office under the Animals (Scientific Procedures) Act 1986.

### Sample preparation

Samples were prepared as described in ^5^ and ^28^. In brief, mice were sacrificed and 600 µm-thick slabs of the olfactory bulb were sliced in ice-cold dissecting buffer using a Leica VT1200 vibratome and immediately transferred to ice-cold fixative (2% glutaraldehyde in 150 mM sodium cacodylate buffer pH 7.4, 300 mOsm/l). Next morning, fixative was washed and the OB slab was then imaged at a 2-photon microscope (^29^, Scientifica Multiphoton VivoScope, coupled with a SpectraPhysics MaiTai DeepSee laser tuned to 940 nm). Slabs were then stained with heavy metals using an established ROTO protocol^5,28^. Subsequently, samples were dehydrated with increasing ethanol solutions (75%, 90%, 2×100%), transferred to propylene oxide, and infiltrated with hard epon mixed with propylene oxide in increasing concentrations (25%, 50%, 75%, 2×100%). Finally, samples were polymerised individually into plastic moulds for 72 h at 70°C.

### Lab X-ray CT imaging

After staining and embedding, the samples were imaged with a laboratory-source X-ray microscope (LXRT; Zeiss Xradia 510 Versa, 40 kV, 3 W, LE2-4 filter, 1601 projections over 180° of sample rotation, 3–20 s exposure time per frame, one tile, 3–6 μm pixel size). This yielded the X-ray volume datasets used to target the fs laser ablation.

### Femtosecond laser milling

The samples were mounted to standard SEM aluminium stubs using conductive silver paint, then sputter coated with palladium to facilitate the SEM imaging of the samples for targeting of the laser. The coating does not interfere with the imaging techniques subsequent to the pillar cutting, as it remains only on the top surface of the pillars. Initially, optimum laser parameters for pillar milling in these samples were determined on resin blocks (with no embedded tissue) to be laser power 20%, pulse frequency 100 kHz, scan speed 500 mm/s, and a spiral hatching pattern in outside-in direction repeated 75 to 100 times, depending on the sample thickness. The milling depth and, thus, the pillar height was adjusted by the number of repetitions of that milling process - up to 5 times with the focus plane being lowered by 0.2 mm each time to adapt the laser focus to the increasing depth.

To perform the targeted pillar milling on a sample imaged by lab X-ray CT, the sample stub was mounted on a custom sample holder and loaded into the main chamber of a ZEISS Crossbeam 350 FIB-SEM instrument with a fs laser (TRUMPF SE + Co. KG, Ditzingen, Germany, wavelength 515 nm, pulse length <350 fs, pulse repetition rate of 1 kHz to 1 MHz, focus spot diameter <15 µm) coupled into a processing chamber attached to the instrument’s airlock. Conducting the laser ablation in a separate processing chamber avoids contamination of the instrument’s main chamber by debris produced by the laser ablation. The sample holder enables milling target coordinate transfer between the SEM and the laser by registering four reference marks on the holder both in the SEM and in the laser reference frames ^30^.

A low magnification SEM image of the sample was acquired with the ZEISS Atlas software on the Crossbeam, and the X-ray volume dataset of the sample was loaded into the same workspace in that software. The X-ray dataset was aligned with and matched to the SEM image on three distinct points visible both in the SEM image and in the part of the X-ray volume that represents the sample surface. Target locations for pillar milling were identified by scrolling through the X-ray volume until the ROIs were encountered. Laser patterns for pillar milling were placed in the SEM image, exactly above the identified ROI positions. Then, the correlation between the SEM coordinate system and the laser coordinate system was established by performing the sample holder registration procedure mentioned above. Finally, the pillar target coordinates were transferred to the laser coordinate system, and the laser milling was executed in the laser processing chamber.

### Sample mounting

Following fs laser milling, pillars were separated from the base using a hand-held single-edge blade and a wooden toothpick with a small trace of cyanoacrylate glue at the tip. The pillars were then individually mounted on aluminium pins suitable for the subsequent imaging technique (either SEM or 3View2-compatible pins were used successfully). The pillars were glued to the center of the pin platform with a small trace of cyanoacrylate glue. For synchrotron imaging, this setup sufficed. For SBF-SEM imaging, the mounted pins were coated on the sides with silver epoxy paint (SPI supplies) and sputter coated with 5 nm of palladium. For FIB-SEM imaging, the unpainted pin was sputter coated with 30 nm of platinum. Individual pins could be manipulated under various magnifying lens or stereoscope setups. Representative images of the mounted pillars were obtained with a Leica M205C stereoscope equipped with a Leica DMC4500 camera.

### Synchrotron CT imaging

Synchrotron imaging was performed at the ID19 microtomography beamline at ESRF as described before^5^. This imaging modality benefits from the cylindrical geometry of the sample. It employs propagation-based phase contrast (26 keV, 1800 projections over 180° of sample rotation, 0.1 s exposure time per frame, LuAG:Ce 25 µm scintillator, pco.edge 5.5 camera with an effective pixel size of 650 nm, 10x objective lens/0.28 numerical aperture).

### FIB-SEM imaging

FIB-SEM data was collected using a Crossbeam 540 FIB-SEM with Atlas 5 for three-dimensional tomography acquisition (Zeiss, Cambridge). The aluminium pin with the Pt coated pillar attached was further mounted on a 12.7 mm SEM stub; the stub was trimmed to have one straight edge and the pin was mounted perpendicular to this edge using carbon cement. In doing so, it was possible to access both the top face and side of the pillar for ion beam milling. An additional 30 nm platinum coat was then applied. Milling was performed at an accelerating voltage of 30 kV with a combination of currents ranging from 65 nA for coarse milling to 700 pA for surface polishing. During image acquisition, the SEM was operated at an accelerating voltage of 1.5 kV with 1.5 nA current. Electron micrographs were acquired at a variety of per-pixel resolutions using a 10 or 20 μs dwell time.

### SBF-SEM imaging

SBF-SEM imaging was performed as described previously ^5,27^. Volumes were acquired on a 3View2-Zeiss Merlin SBF-SEM under high vacuum (1.5 kV, 0.5 nA, 0.5 μs/pixel, 10 nm pixels, resulting in a surface dose of 15.6 e^-^/nm^2^) using an OnPoint (Gatan) back-scattered electron detector. Nominal cutting thickness was 50 nm. The dataset contains 100 sequential images each covering 4096*4096 pixels, without any slice loss. The dataset was registered using Voxelytics Align (scalable minds, Potsdam, Germany), which uses pairwise feature matching between adjacent slices, RANSAC inlier detection and global relaxation to create mesh-tessellated affine transformations. The resulting dataset maps a volume of 42.4 * 41.4 * 5 µm^3^ with 10*10*50 nm^3^ voxels. All datasets were processed and managed using a webKnossos data storage, display and annotation infrastructure^31^.

### Correlative multimodal imaging

Datasets from 2-photon and lab and synchrotron X-ray tomography were acquired from the same specimen at targeted locations and warped to a common space as previously described^5^. Briefly, datasets obtained with different imaging modalities that mapped the same sample volume could be warped to a common space by manually logging conserved pairs of landmarks using BigWarp^32^. The nature and number of landmark pairs necessary depended on the imaging modalities being paired. *Ex vivo* 2-photon to LXRT could be warped relying on major histological features such as tissue boundaries and glomerular contours. LXRT to SXRT could be warped relying on these and finer features, including large blood vessel branching points or large cell nuclei. To target specific locations across datasets, a warp graph was built containing all related datasets (nodes) and warp functions across dataset pairs (edges) using previously described custom code^33^. This allowed easily warping any annotation (for example the location of a fluorescent glomerulus in the 2-photon dataset) across any of the linked datasets in the graph (such as the LXRT dataset).

